# Bioinformatic Surveillance Leads to Discovery of Two Novel Putative Bunyaviruses Associated with Black Soldier Fly

**DOI:** 10.1101/2023.06.23.545759

**Authors:** Hunter K. Walt, Emilia Kooienga, Jonathan A. Cammack, Jeffery K. Tomberlin, Heather Jordan, Florencia Meyer, Federico G. Hoffmann

## Abstract

The black soldier fly (*Hermetia illucens,* BSF) has emerged as an industrial insect of high promise because of its ability to convert organic waste into nutritious feedstock, making it an environmentally sustainable alternative protein source. As global interest rises, rearing efforts are also upscaled, which is highly conducive to pathogen transmission. Viral epidemics have stifled mass-rearing efforts of other insects of economic importance, such as crickets, silkworms, and honeybees, but little is known about the viruses that associate with BSF. Although it is thought that BSF are unusually resistant to pathogens because of their expansive antimicrobial gene repertoire, surveillance techniques could be useful to identify emerging pathogens and common BSF microbes. In this study, we used high-throughput sequencing data to survey BSF larvae and frass samples, and we identified two novel bunyavirus-like sequences. Our phylogenetic analyses grouped one in the family *Nairoviridae*, and the other with two unclassified bunyaviruses. We describe these putative novel viruses as BSF Nairovirus-like 1 and BSF uncharacterized bunyavirus-like 1. We identified candidate segments for the full BSF Nairovirus-like 1 genome using a technique based on transcript co-occurrence, and only a partial genome for BSF uncharacterized bunyavirus-like 1. These results emphasize the value of routine BSF colony surveillance and add to the number of viruses associated with BSF.

## INTRODUCTION

Insects have become a point of interest as a sustainable alternative source of food and feed. Insect rearing is less environmentally impactful than traditional food and feed production, but to produce a substantial amount of biomass, insects must be reared in mass (van Huis, 2020). Mass rearing of animals is highly conducive to pathogen transmission as many individuals are reared in close quarters. Insects are no exception, and the production of insects of economic importance such as crickets (Orthoptera: *Gryllidae*), silkworms (Lepidoptera: *Bombycidae*), and honeybees (Hymenoptera: *Apidae*) has been historically stifled by viral epidemics (Reviewed by Eilenberg *et al*. 2015).

The black soldier fly (BSF) (*Hermetia illucens*) is an insect of emerging industrial importance. This is because of its ability to convert organic waste to highly nutritious biomass for animal feed and potentially food for humans (Tomberlin and van Huis, 2020). It has also been investigated as a source of antimicrobial compounds, antioxidants, and biodiesel (Zhu *et al*. 2019, Xia *et al*. 2021, Lu *et al*. 2022). As a result, there is global interest in upscaling rearing of the species; however, relatively little is known about the microbes that could help or hinder its production (Joosten *et al*. 2020). Furthermore, it has been proposed that BSF are unusually resistant to pathogens because of their large repertoire of antimicrobial peptides (Park *et al*. 2014, Zhan *et al*. 2020, Moretta *et al*. 2020). Until recently, no known viruses associated with BSF had been identified. Pienaar *et al*. (2022) mined publicly available BSF data for endogenous viral elements (EVEs) and viruses and found multiple EVEs and one exogenous toti-like virus. This suggests that there is a history of BSF-virus interactions and emphasizes the need for investigating BSF-virus interaction.

In this study, we use a computational approach to survey RNA-seq datasets for viruses associated with BSF. We use transcriptomic data generated from previous BSF studies to search for putative novel viral-like sequences associated with BSF. We use a sequence similarity-based approach to identify putative viral sequences in our data, infer phylogenetic relationships based on the RNA-dependent RNA polymerase (RdRp) gene, and identify candidate sequences of diverged genome segments of these viruses based on co-occurrence of transcripts between samples (Batson *et al*. 2021). Our results emphasize the potential of high-throughput sequencing technology as a biological surveillance tool that can also identify emerging pathogens of BSF. In addition, this type of data provides foundations to develop molecular diagnostic tools to actively monitor colonies for viral infection (Semberg *et al*. 2019).

## MATERIALS AND METHODS

### Sample Preparation and Sequencing

BSF larval gut and frass reads were generated from industrial scale experiments. Briefly, *Arthrobacter AK19 or Rhodococcus rhodochrous* 21198 were used along with spent brewer’s grain, a waste stream that would be reasonably used as an industrial source of BSF larval diet. 8.0 g of either *Arthrobacter AK19* or *R. rhodochrous 21198* were added to 6.0 kg of spent brewer’s grain diet per pan. The control diet consisted of spent brewer’s grain only. Approximately 10,000 larvae were added to each pan of either non-supplemented or supplemented treatments. Each treatment condition (*Arthrobacte*r-supplemented, *Rhodococcus*-supplemented, and control) had four replicates (or pans). The larvae were allowed to feed constantly on the initially placed diet. At designated timepoints of the 10-day experiment, a subset of larvae was removed from the pans, weighed, and frozen for later analysis. A sample of the initial spent grain, as well as insect digestate-substrate (hereafter referred to as frass) was also collected (at corresponding timepoints to larval collection) and frozen for later analysis.

Larvae were surface sterilized by dipping in 10% bleach solution for 1 minute, followed by two successive deionized water washes for one minute. Following this, the intestinal tract (gut) was dissected from the larvae. RNA was purified from larval guts, frass, and the initial spent grain diet using the Direct-zol™ RNA MiniPrep kit, following the manufacturer’s protocol, and RNA products were quantified with a Qubit® 2.0 fluorometer, and then ran on a gel to determine RNA quality. The NEB Ultra RNA Library Kit (New England Biolabs) was used to convert the RNA to cDNA, avoiding steps to remove rRNA, and preserving total RNA concentrations. cDNA libraries were multiplexed using NEBNext Oligos for Illumina to create five samples of pooled libraries. Resulting multiplexed libraries were quality verified using a 4150 TapeStation System (Agilent) then submitted for shotgun whole metatranscriptome sequencing using an Illumina® HiSeq 2000 instrument. Sequences were deposited in the Sequence Read Archive database under BioProject ID: PRJA976287.

### Metatranscriptome Assembly and Virus Discovery

The paired end reads were quality checked using fastQC v.0.11.9 (Andrews, 2010) and quality trimmed using trimmomatic v.0.39 with the options ILLUMINACLIP:TruSeq3-PE-2.fa:2:30:10, SLIDINGWINDOW:4:15, and MINLEN: 30 (Bolger *et al*. 2014). All read libraries were then mapped to the reference BSF genome (GCA_905115235.1) (Generalovic *et al*. 2021) using HISAT2 v.2.2.1 (Daehwan *et al*. 2019), and only the non-BSF reads were retained using the –un-conc option. The non-BSF reads were subsequently assembled using Trinity v2.14.0 (Grabherr *et al*. 2011) through Trinity’s Docker container (Merkel, 2014).

### Identification of Conserved Viral Fragments

To identify putative viral sequences, all transcripts in the assembled transcriptomes were renamed by sample ID and a unique identifier and concatenated. Next, the concatenated transcriptome was clustered using CD-hit-EST (Li and Godzik, 2006) at a threshold of 95% identity, a word size of 5, and a minimum length of 500 nt. The representative sequences from these clusters were then annotated using DIAMOND + BLASTx using the –very-sensitive flag and an e-value threshold of 1e^-2^. Sequences matching RNA viruses were considered of probable viral origin. To eliminate false positives, these sequences were further inspected to establish the presence of ORFs of viral origin using NCBI’s ORFfinder (https://www.ncbi.nlm.nih.gov/orffinder/). All large ORFs were translated and aligned to NCBI’s nr protein database using the BLASTp webserver (https://blast.ncbi.nlm.nih.gov/Blast.cgi). Viral conserved protein domains were verified using NCBI’s conserved domain database (https://www.ncbi.nlm.nih.gov/Structure/cdd/cdd.shtml) and InterProScan (https://www.ebi.ac.uk/interpro/search/sequence/) (Quevillon *et al*. 2005). Sequences matching RNA viruses that passed the criteria listed above were considered as viral conserved sequences.

### Identification of Diverged Viral Fragments

To find diverged genomic segments of viruses in the assembled transcriptomes, we implemented a method to identify transcripts that co-occur with viral conserved sequences (Batson *et al*. 2021). We assumed that diverged genomic segments would not have significant BLAST hits, so we only considered unannotated transcripts in the ensuing analysis. We wrote a python script that determines all the samples where a viral conserved sequence is found. Next, we used two metrics that assess how often any other transcripts co-occur with the viral conserved sequence: a viral co-occurrence metric (*V_co_*) and a transcript co-occurrence metric (*T_co_*). *V_co_* measures how frequently a transcript is found in the same sample as a viral conserved sequence and is used to filter out transcripts that are not frequently found in the sample as the viral conserved sequence. Mathematically, *V_co_* is equal to the cardinal number of the intersection of *V* and *T* divided by the cardinal number of *V*, where *V* is the set of samples where a viral conserved sequence is found, and *T* is the set of samples that a transcript occurs in (**Equation 1**).

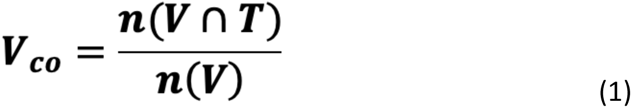

Because *n*(*V*∩*T*) ≤ *n*(*V*), *V_co_* ranges from 0, where the transcript and viral conserved sequence of interest are never found in the same sample, to 1, where the transcript and viral conserved sequence of interest are only found together. Intermediate values of V_co_ (0 < V_co_ < 1) indicate that the viral conserved sequence and the transcript co-occur in some samples, but not in all.

The transcript co-occurrence metric (*T_co_*) is a complementary metric that assesses whether a transcript is found in samples without the viral conserved sequence. *T_co_* is described by **equation 2** and is the cardinal number of the intersection of *V* and *T* divided by the cardinal number of *T*.

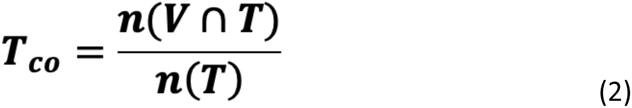

Because *n*(*V*∩*T*) ≤*n*(*T*), *T_co_* ranges from 0, where the transcript and the viral conserved sequence of interest are never found in the same sample, to 1, where the transcript is only found with the viral conserved sequence. Intermediate values of T_co_ (0 < T_co_ < 1) indicate that the transcript is found in samples with the viral conserved sequence, but also in additional samples. Ideally, *T_co_* and *V_co_* will be close to 1. In our case, we used a threshold of *V_co_* = 0.75 and *T_co_* = 0.5 to designate a transcript as putatively viral. The remaining transcripts were filtered by an ORF size > 500 nucleotides and inspected for other viral attributes using NCBI’s conserved domain database, BLAST, and InterProscan.

### Phylogenetic Analysis

To determine the evolutionary affinities of the novel viruses, we performed a phylogenetic analysis using the RNA-dependent RNA-polymerase (RdRp) gene from a selection of bunyaviruses. The amino acid sequence for each bunyavirus RdRp was downloaded from NCBI (https://www.ncbi.nlm.nih.gov/). The ORF containing the novel viruses’ RdRp was extracted and translated using NCBI’s ORFfinder. All resulting amino acid sequences were aligned using mafft v7.490 using the E-INS-I algorithm and the –maxiterate option set to 1000 (Kathoh and Standley 2013). ModelFinder (Kalyaanamoorthy *et al*. 2017) was used through the IQ-TREE2 v2.0.7 to find the best evolutionary model, and IQ-TREE2 v2.0.7 was used to infer the most likely tree (Minh *et al*. 2020). Branch support was measured using ultrafast bootstrap with 1000 replicates, the Shimodaira-Hasegawa-like approximate likelihood ratio test (SH-aLRT) with 1000 replicates, and the aBayes test (Anisomova *et al*. 2011, Nguyen *et al*. 2015, Minh *et al*. 2013).

### Public Data Analysis

All publicly available BSF transcriptomic data (n=116, **Supplementary Table 3**) was downloaded using the SRA toolkit v.3.0.0 (https://hpc.nih.gov/apps/sratoolkit.html) and mapped to the candidate viral segments using BWA-mem v2.2.1 (Vasimuddin *et al*. 2019). Resulting BAMs were sorted and indexed using SAMtools v.1.6 (Li et al. 2009). The presence of viral transcripts in public data was inspected manually using Integrative Genomics Viewer v.2.14.1 (Robinson *et al*. 2011). If a putative viral transcript was found, the corresponding SRA dataset was processed using the same pipeline described earlier in the methods, added to the concatenated metatranscriptome, and the co-occurrence analysis was run again.

## RESULTS

We mapped 13,352,390 reads generated from 38 BSF larvae and frass samples to the BSF reference genome (GCA_905115235.1) (Generalovic *et al*. 2021). An overview of the samples and the percentage of the reads that mapped to the BSF genome is shown in **Table 1**. For each sample, we assembled the reads that did not map to the BSF genome, resulting in 38 metatranscriptomes. CD-hit clustering of the metatranscriptomes resulted in 7,962 distinct sequence clusters, all of which shared greater than 95% sequence identity. The representative sequence from each cluster was annotated using diamond BLASTx, resulting in 7,830 (98.3%) annotations. We filtered these results for viral hits and found two transcripts with significant hits to RdRps of arthropod-infecting bunyaviruses: one transcript was 5,696 nucleotides in length and the other was 4,542 nucleotides. We hereon refer to the virus with the longer RdRp transcript as BSF uncharacterized bunyavirus-like 1, and the other we refer to as BSF Nairovirus-like virus 1. Both transcripts shared their top five BLAST hits (**Table 2**). We confirmed the presence of the bunyaviral RdRp domain using the NCBI conserved domain database and InterProscan. We found the BSF uncharacterized bunyavirus-like 1 encodes for a bunyavirus-like RdRp conserved domain between nucleotides 3,950-4,294 (E-value = 2.43e^-6^), and BSF Nairovirus-like 1 encodes for a bunyavirus-like RdRp conserved domain between nucleotides 2,052-2,888 (E-value = 1.10e^-15^). Other putative viral sequences such as RNA phages were found and were not included in our analyses because they are likely associated with microbes (Callanan et al. 2018) (**Supplementary Table 1**).

**Table 1:** Sample Overview. A summary of each sample, number of reads, percent mapped to BSF genome, sample type, time point, and treatment.

| Sample | Number of Reads | Alignment Rate | Sample Type | Timepoint | Treatment |
| --- | --- | --- | --- | --- | --- |
| SG1 | 35728 | 0.59% | Spent Grain | 0 | Control |
| T3LA2 | 23447 | 32.26% | Larvae | 3 | *Arthrobacter* |
| T3LA4 | 1017935 | 85.79% | Larvae | 3 | *Arthrobacter* |
| T3LC1 | 18192 | 15.88% | Larvae | 3 | Control |
| T3LC2 | 22027 | 38.51% | Larvae | 3 | Control |
| T3LC4 | 162829 | 86.39% | Larvae | 3 | Control |
| T3LR1 | 1137578 | 85.40% | Larvae | 3 | *Rhodococcus* |
| T3LR3 | 12247 | 24.13% | Larvae | 3 | *Rhodococcus* |
| T3WA1 | 604 | 1.90% | Frass | 3 | *Arthrobacter* |
| T3WA2 | 7373 | 2.10% | Frass | 3 | *Arthrobacter* |
| T3WA4 | 7613 | 1.83% | Frass | 3 | *Rhodococcus* |
| T3WC1 | 17569 | 0.73% | Frass | 3 | Control |
| T3WC2 | 21706 | 1.21% | Frass | 3 | Control |
| T3WC4 | 52404 | 0.53% | Frass | 3 | Control |
| T3WR1 | 315689 | 0.90% | Frass | 3 | *Rhodococcus* |
| T3WR3 | 47411 | 0.47% | Frass | 3 | *Rhodococcus* |
| T6LA1 | 388220 | 86.20% | Larvae | 6 | *Arthrobacter* |
| T6LA2 | 1264636 | 90.31% | Larvae | 6 | *Arthrobacter* |
| T6LA3 | 212178 | 77.46% | Larvae | 6 | *Arthrobacter* |
| T6LA4 | 125 | 9.20% | Larvae | 6 | *Arthrobacter* |
| T6LC1 | 89711 | 60.09% | Larvae | 6 | Control |
| T6LC2 | 551 | 26.23% | Larvae | 6 | Control |
| T6LC3 | 1500632 | 86.55% | Larvae | 6 | Control |
| T6LC4 | 833147 | 93.56% | Larvae | 6 | Control |
| T6LR1 | 66 | 0.76% | Larvae | 6 | *Rhodococcus* |
| T6LR2 | 210 | 30.71% | Larvae | 6 | *Rhodococcus* |
| T6LR3 | 75 | 53.33% | Larvae | 6 | *Rhodococcus* |
| T6LR4 | 456745 | 95.04% | Larvae | 6 | *Rhodococcus* |
| T6WA1 | 995664 | 5.14% | Frass | 6 | *Arthrobacter* |
| T6WA2 | 956959 | 2.84% | Frass | 6 | *Arthrobacter* |
| T6WA4 | 10006 | 1.82% | Frass | 6 | *Arthrobacter* |
| T6WC1 | 684749 | 2.26% | Frass | 6 | Control |
| T6WC2 | 737122 | 3.18% | Frass | 6 | Control |
| **T6WC3** | 93147 | 15.04% | Frass | 6 | Control |
| **T6WC4** | 79076 | 1.98% | Frass | 6 | Control |
| **T6WR1** | 20437 | 1.34% | Frass | 6 | *Rhodococcus* |
| **T6WR2** | 1130727 | 2.12% | Frass | 6 | *Rhodococcus* |
| **T6WR3** | 997855 | 0.06% | Frass | 6 | *Rhodococcus* |

**Table 2:**
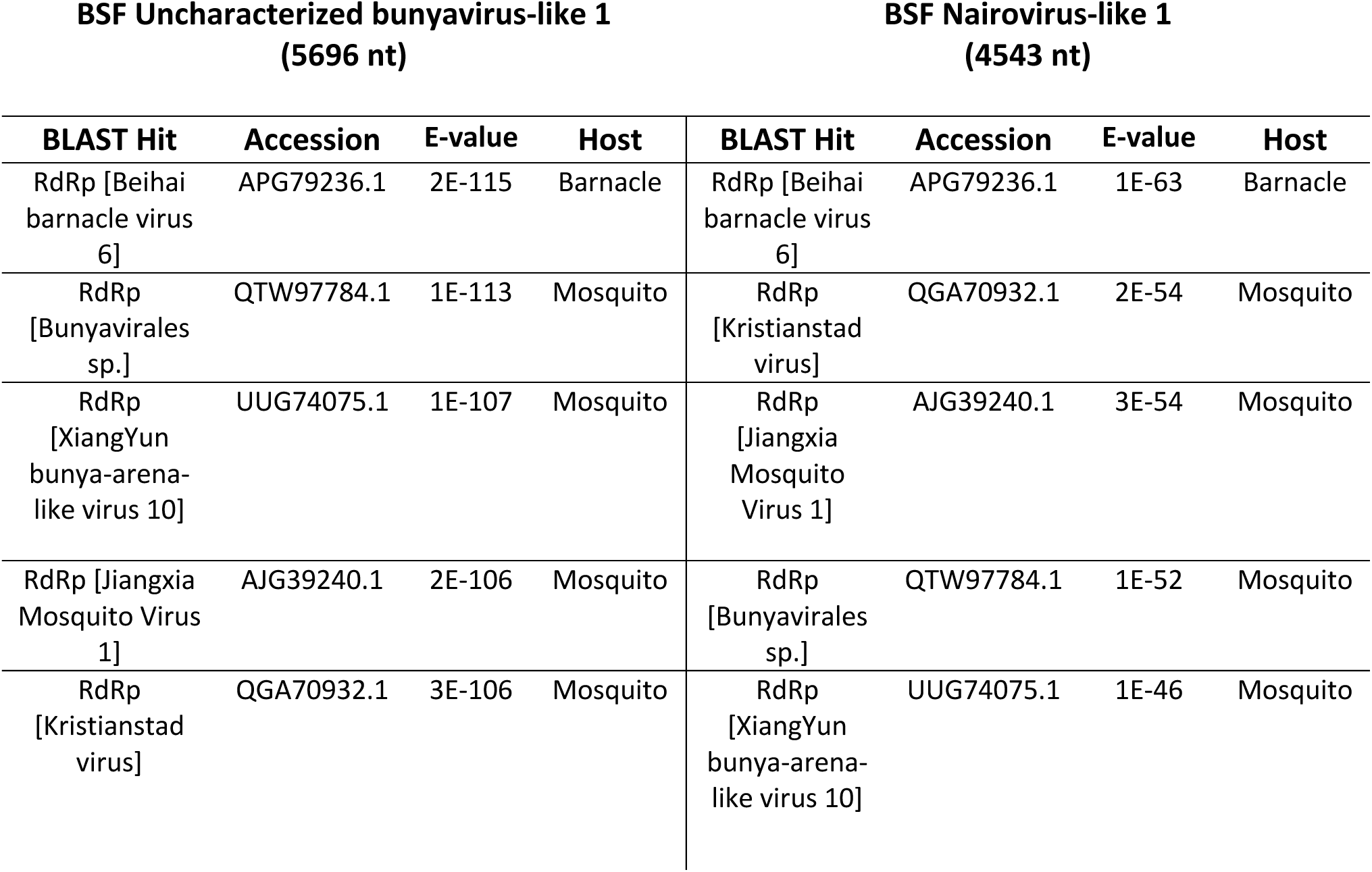
Top five BLAST hits of the two putative novel bunyavirus L segments discovered in this study. The Table contains Each BLAST hit, the subject’s accession number, the corresponding e-value, and the host from which the virus was identified. Notably, all hosts are arthropods.

| BSF Uncharacterized bunyavirus-like 1<br>(5696 nt) |  |  |  | BSF Nairovirus-like 1<br>(4543 nt) |  |  |  |
| --- | --- | --- | --- | --- | --- | --- | --- |
| BLAST Hit | Accession | E-value | Host | BLAST Hit | Accession | E-value | Host |
| RdRp [Beihai barnacle virus 6] | APG79236.1 | 2E-115 | Barnacle | RdRp [Beihai barnacle virus 6] | APG79236.1 | 1E-63 | Barnacle |
| RdRp [Bunyavirales sp.] | QTW97784.1 | 1E-113 | Mosquito | RdRp [Kristianstad virus] | QGA70932.1 | 2E-54 | Mosquito |
| RdRp [XiangYun bunya-arena-like virus 10] | UUG74075.1 | 1E-107 | Mosquito | RdRp [Jiangxia Mosquito Virus 1] | AJG39240.1 | 3E-54 | Mosquito |
| RdRp [Jiangxia Mosquito Virus 1] | AJG39240.1 | 2E-106 | Mosquito | RdRp [Bunyavirales sp.] | QTW97784.1 | 1E-52 | Mosquito |
| RdRp [Kristianstad virus] | QGA70932.1 | 3E-106 | Mosquito | RdRp [XiangYun bunya-arena-like virus 10] | UUG74075.1 | 1E-46 | Mosquito |

To phylogenetically analyze the two novel putative viruses, we translated their coding sequences and aligned them with 55 known bunyavirus RdRp amino acid sequences retrieved from NCBI to construct a phylogenetic tree using the VT+F+R7 amino acid substitution model, which was chosen as the best fit by ModelFinder (**Figure 1**). The sequences were chosen to represent all bunyavirus families documented in NCBI (Wang *et al*. 2022). The phylogeny shows BSF uncharacterized bunyavirus-like 1 groups with two taxonomically uncharacterized arthropod-infecting bunyaviruses: Kristianstad virus (detected in mosquitoes) (Pettersson *et al*. 2019) and Beihai barnacle virus 6 (detected in barnacles) (Shi *et al*. 2016). Interestingly, BSF Nairovirus-like virus 1 groups with the RdRp of Abu Hammad virus, which is in the *Nairoviridae* family that was isolated from a tick. Many of the viruses in the *Nairoviridae* family also infect vertebrates and can be transmitted by arthropod vectors (Walker et al. 2016). The full phylogenetic tree with no collapsed nodes is shown in **supplementary figure 1.**

**Figure 1:**
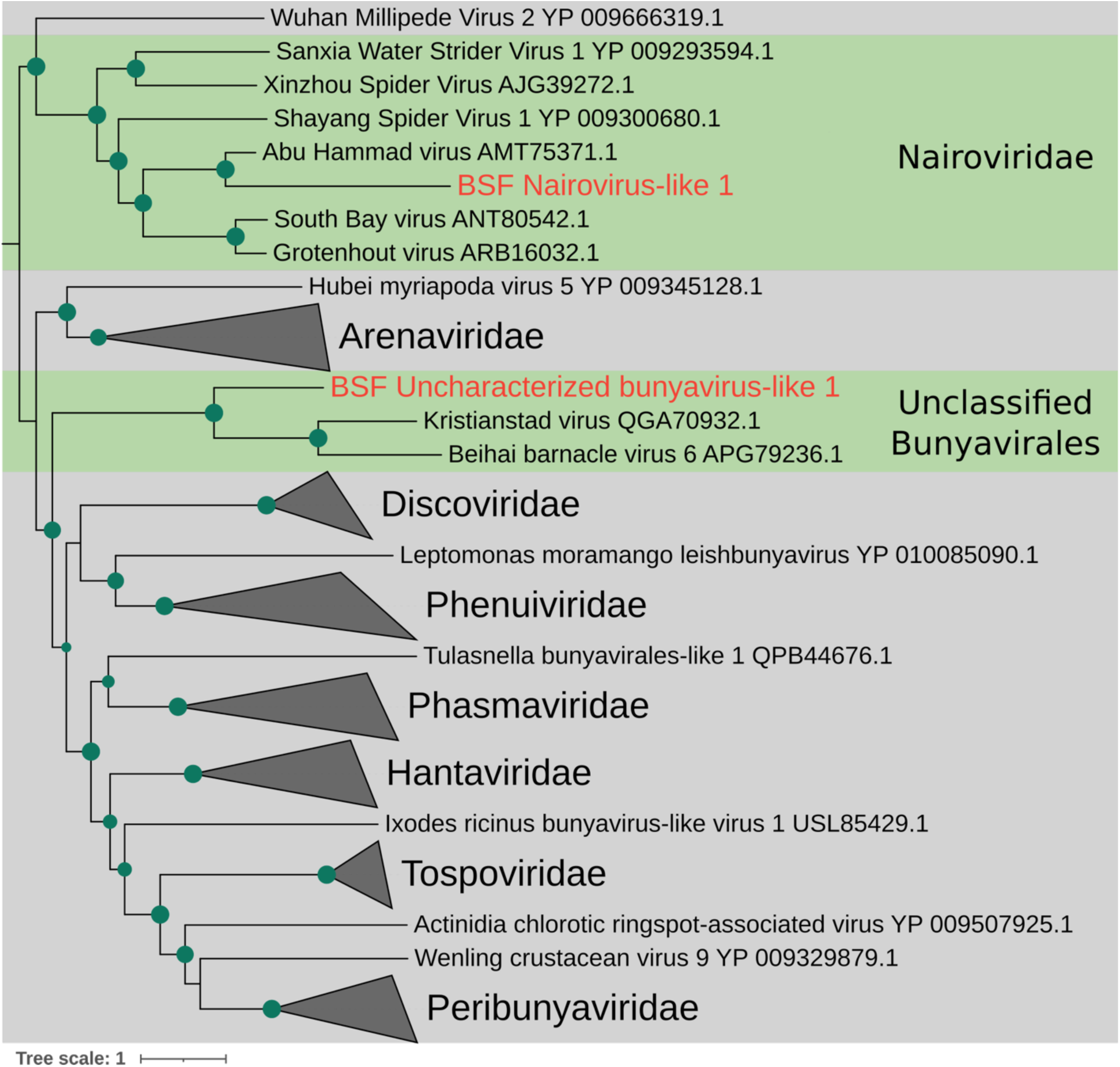
Phylogenetic tree of Bunyavirus RNA-dependent RNA polymerase. The viruses discovered in this paper are in red text, and their clade is highlighted in green. The green dots along the branches represent a minimum bootstrap value of 75. Families that did not contain the novel putative viral transcripts were collapsed at their common ancestral node. The tree was rooted at the midpoint.

**Figure 2:**
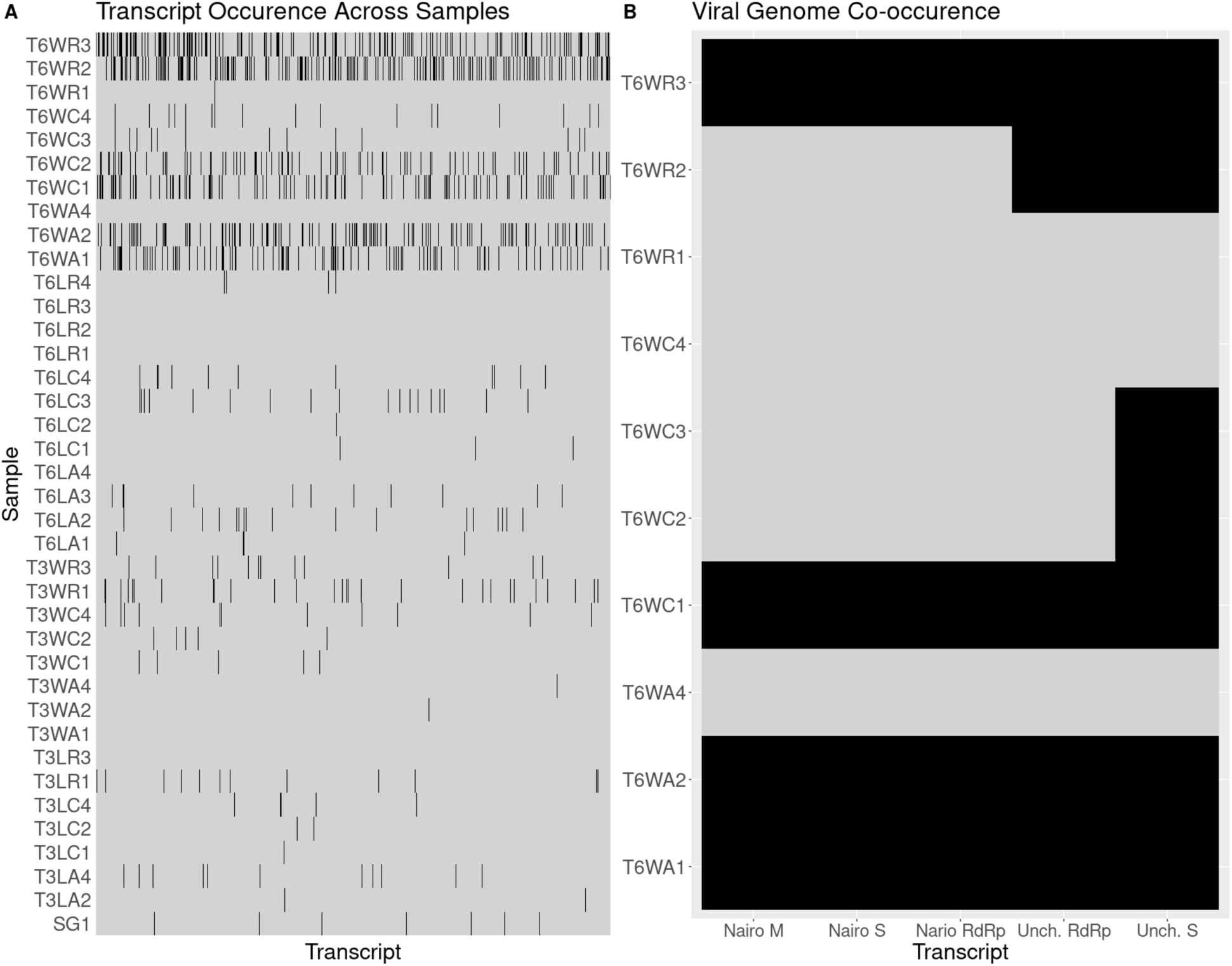
Transcript co-occurrence across samples. A.) Occurrence matrix of all CD-hit transcript clusters across samples, where black indicates presence and grey indicates absence. The X axis is all transcript clusters from CD-hit at 95% sequence identity. B.) Occurrence matrix only using the putative viral sequences discovered in this study.

Generally, bunyaviruses have a segmented genomic architecture with a large (L) segment containing the conserved RdRp domain, a medium (M) segment, and a small (S) segment. Because of the error-prone replication by RdRp and extreme selection pressures from host-virus interactions, the M and S segments of bunyaviruses can become very diverged and sometimes impossible to detect based on sequence similarity. We did not find any BLAST results that matched other viral M or S segments, so we employed an approach based on transcript co-occurrence across samples, similar to the approach taken by Batson *et al*. (2021). We assumed that if an L segment occurred in multiple samples, the M and S segments would also occur in those samples. Because BSF uncharacterized bunyavirus-like 1 occurred in five samples (out of 38), and BSF Nairovirus-like 1 occurred in four samples (out of 38), we were able to employ this method.

We wrote a python script that takes a defined list of viral conserved sequences (the two novel bunyavirus RdRp transcripts in our case), determines the samples where the viral conserved sequence occurred, and returns a viral co-occurrence metric (*V_co_*, **equation 1**) and a transcript co-occurrence metric (*T_co_*, **equation 2**). *V_co_* is a metric to ensure that a transcript occurs in the same samples as the viral conserved sequence, while *T_co_* is a metric to assess whether a transcript co-occurs with the viral conserved sequence and no other samples. Ideally, *V_co_* and *T_co_* should equal 1 (i.e., a transcript perfectly occurs with a viral conserved sequence and only in those samples), but because of limited sequencing depth, we set the *V_co_* threshold to 0.75, and the *T_co_* threshold to 0.5 to identify candidate M and S segments. We assumed that diverged viral genome segments had no blast hits with an e-value higher than our 1e^-2^ threshold, so only unannotated transcripts were considered. We found 5 candidate representative transcripts for M or S segments for uncharacterized BSF bunyavirus-like 1, and 7 candidate transcripts for BSF Nairovirus-like 1. These transcripts were filtered by the presence of complete ORFs greater than 500 nucleotides in length, as bunyaviral M and S segments typically have continuous ORFs throughout their genomic segments. This left us with four potential candidates for the diverged M and S segments, but all four of these transcripts met the co-occurrence requirements for both viruses.

To determine which candidate M and S segments go with each virus, we mined all available BSF transcriptomic data by mapping their reads to the putative L segments. No reads mapped to BSF Nairovirus-like 1, but reads from six datasets mapped to BSF uncharacterized bunyavirus-like 1 (**Supplementary Table 2**). Notably, all the public data this virus occurred in were processed in the same lab as the samples used in this study, but for a different experiment (BioProject Accession: PRJNA542977). The datasets containing BSF uncharacterized bunyavirus-like 1 were processed in the same way as the reads generated in our study (i.e., trimmed with Trimmomatic, quality checked with FastQC, and assembled with Trinity), and the transcripts were added to the co-occurrence analysis. After repeating the co-occurrence analysis with the SRA data included, BSF uncharacterized bunyavirus-like 1 only had one candidate that met our criteria. This sequence was also present in the first co-occurrence analysis. Two candidate diverged viral segments appeared with BSF Nairovirus-like 1 with *V_co_* = 1 and *T_co_* = 1. Thus, we assigned these two transcripts as the M and S segments for BSF Nairovirus-like 1, which are 2,837 and 922 nucleotides long, respectively (**Figure 3A**). We assigned the transcript identified with BSF uncharacterized bunyavirus-like 1 as its S segment, as it is 853 nucleotides long (**Figure 3B**).

**Figure 3:**
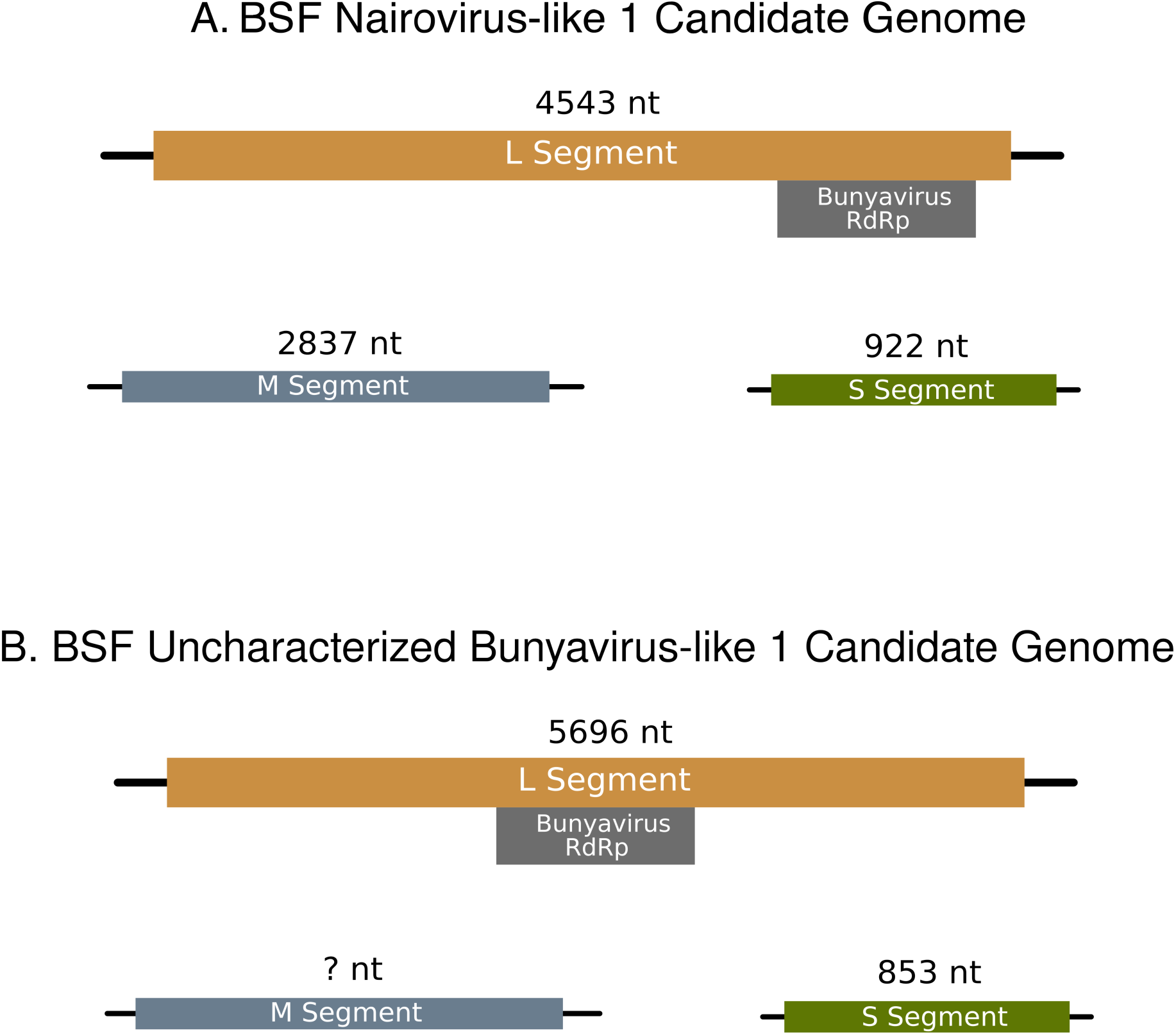
Genome Segments of the two bunyaviruses discovered in this study. A.) The proposed genome of BSF Nairovirus-like 1. The L, M, and S segments are 4,543, 2,837, and 922 nucleotides, respectively. The bunyavirus RdRp conserved domain lies between nucleotides 3,950-4,294 (E-value = 2.43e^-6^) on the L segment. **B.)** The proposed genome of BSF Uncharacterized Bunyavirus-like 1. The L and S segments are 5696 and 853 nucleotides, respectively. No transcript met the requirements for an M segment candidate. The bunyavirus RdRp conserved domain lies between nucleotides 2,052-2,888 (E-value = 1.10e^-15^) on the L segment.

## DISCUSSION

In this study, we discovered two novel putative bunyaviruses associated with BSF. To our knowledge, they are only the second and third viruses associated with BSF. Although we cannot confirm if these viruses are pathogenic to BSF, diagnostic molecular tools can be used to investigate if these viruses coincide with BSF disease symptoms. We show that both viruses share most recent common ancestors with other insect viruses, although both viruses were only found in frass samples. We suspect two reasons why these viruses were only found in frass samples. 1.) The virus might infect tissues other than the gut, which are the only BSF tissues used in this study. 2.) Sequencing depth might not have been sufficient to detect these viruses in larval tissues. Alternatively, these viruses may infect components of the BSF frass, as bunyaviruses are diverse and can infect a broad range of hosts (Abudurexiti et al. 2019). Either way, our phylogeny suggests that these are arthropod infecting viruses as their closest phylogenetic relatives also infect arthropods. Interestingly, BSF Nairovirus-like 1 is phylogenetically related to viruses that infect vertebrates, which could be a point of concern for food and feed safety. Future studies should be conducted to confirm the host specificity and pathogenicity of these viruses to ensure that they do not affect BSF rearing. Additionally, we detected BSF uncharacterized bunyavirus-like 1 in 11 datasets (five from this study, six from a previous study, BioProject Accession: PRJNA542977, **Supplementary Table 2**). These datasets were all produced in a single lab, though from different BSF experiments conducted in differing years and in different rearing locations. However, the BSF utilized for each of these experiments were from the same original colony. It will therefore be important to understand the broader distribution of this virus across different BSF strains.

The main limitations of this study relate to the data used, as our sequencing data had poor depth. This may be why we were unable to get a *V_co_* and *T_co_* of one for BSF uncharacterized bunyavirus-like 1, hence why we could not identify a candidate M segment. On the other hand, we show that novel viruses can be discovered even at low sequencing depths, especially when frass samples are used. Finally, this method can be easily applied to any system, although it is optimal to have a high-quality genome to reduce the amount of host reads.

## ACKNOWLEDGEMENTS

We would like to thank Amber E. McDonald for her input on figure design.

## AUTHOR CONTRIBUTIONS

Conceptualization-F.M., H.R.J., F.G.H., H.K.W. J.K.T., J.A.C.; Methodology-H.K.W. and F.G.H.; Supervision-F.M., F.G.H., and H.R.J.; Formal analysis-H.K.W.; Investigation-E.K., H.R.J.; Software-H.K.W.; Writing-original Draft-H.K.W.; Writing-review and editing: F.M., F.G.H. H.R.J., H.K.W., J.K.T., J.A.C.;

## DATA AVAILABILITY STATEMENT

All reads generated from this study were deposited in the NCBI sequence read archive under BioProject ID: PRJNA976287. Python code for co-occurrence analysis can be found at https://github.com/hunterkwalt/virus_co-occur.

