## Supplemental Material for "Bioinformatic Surveillance Leads to Discovery of Two Novel Putative Bunyaviruses Associated with Black Soldier Fly"

**SUPPLEMENTARY MATERIALS:**

**Supplementary Figure 1: Fully uncollapsed bunyavirus phylogenetic tree of the Bunyavirus RdRps used in this study.** This figure is the same tree shown in Figure 1, but all nodes are uncollapsed. All bootstrap values are shown.

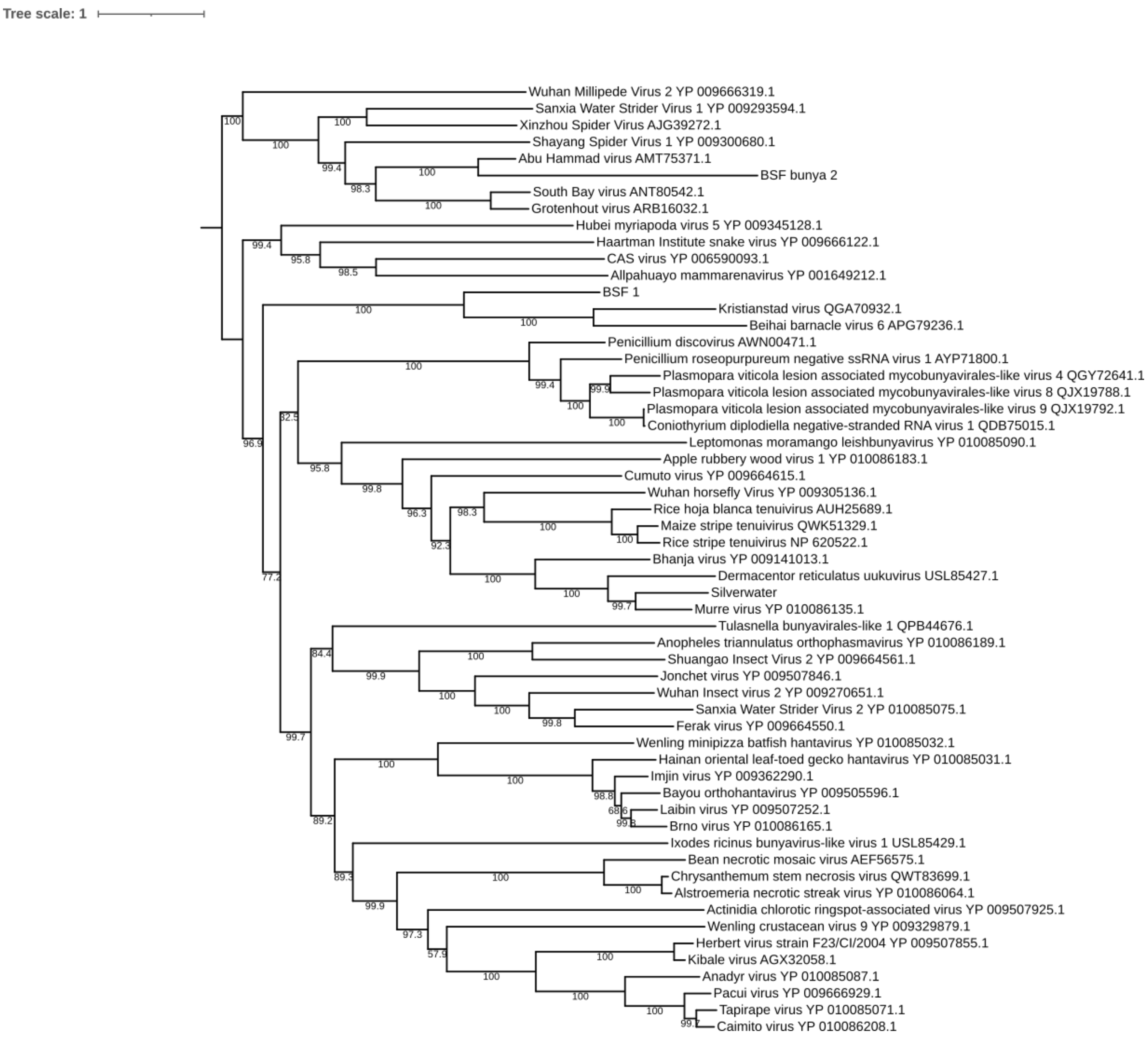

**Supplementary Table 1:** Putative microbe-associated viruses found within BSF. The table only contains contigs with a length greater than 2000.

| Transcript | Length (nt) | Closest BLAST Hit | evalue | Putative Type |
|---|---|---|---|---|
| **T6WA2_S51_DN382_c0_g2_i1** | **3554** | YP_010769592.1: ssRNA phage SRR6960799_20 | **0.0** | **Levivirus (+ssRNA)** |
| **T6WR2_S83_DN3594_c0_g1_i1** | **3472** | URG16190.1: **Leviviridae sp.** | **0.0** | **Levivirus (+ssRNA)** |

**Supplementary Table 2: SRA data that were positive for BSF uncharacterized bunyavirus-like 1.** The table contains the SRA accession number along with the type of sample the virus RdRp transcript was found in.

| SRA Accession | Sample Type |
|---|---|
| **SRR9068902** | **BSF Frass** |
| **SRR9068904** | **BSF Frass** |
| **SRR9068905** | **BSF Frass** |
| **SRR9068906** | **BSF Frass** |
| **SRR9068923** | **Starved BSF Larvae** |
| **SRR9068924** | **Starved BSF Larvae** |

**Supplementary Table 3: List of all SRA data used in this study by accession number.**

**ERR1801985**
**ERR1801986**
**ERR1801987**
**ERR1801988**
**ERR1801989**
**ERR1801990**
**ERR1801991**
**ERR1801992**
**ERR1801993**
**ERR1801994**
**ERR1801995**
**ERR1801996**
**ERR1801997**
**ERR1801998**
**SRR10158821**
**SRR10233312**
**SRR14339782**
**SRR14339783**
**SRR14339784**
**SRR14339785**
**SRR14339786**
**SRR14339787**
**SRR14339788**
**SRR14339789**
**SRR14339790**
**SRR14339791**
**SRR14339792**
**SRR14339793**
**SRR14339794**
**SRR14339795**
**SRR14339796**
**SRR18283674**
**SRR18283675**
**SRR18283676**
**SRR18283677**
**SRR18283678**
**SRR18283679**
**SRR18283680**
**SRR18283681**
**SRR18283682**
**SRR18283683**
**SRR18283684**
**SRR18283685**

| | |
|---|---|
| 61 | **SRR18283686** |
| 62 | **SRR18283687** |
| 63 | **SRR18283688** |
| 64 | **SRR18283689** |
| 65 | **SRR18283690** |
| 66 | **SRR18283691** |
| 67 | **SRR18283692** |
| 68 | **SRR18283693** |
| 69 | **SRR18283694** |
| 70 | **SRR18283695** |
| 71 | **SRR18283696** |
| 72 | **SRR18283697** |
| 73 | **SRR18283698** |
| 74 | **SRR18283699** |
| 75 | **SRR18283700** |
| 76 | **SRR18283701** |
| 77 | **SRR18283702** |
| 78 | **SRR18283703** |
| 79 | **SRR18283704** |
| 80 | **SRR18283705** |
| 81 | **SRR18283706** |
| 82 | **SRR6656085** |
| 83 | **SRR6656086** |
| 84 | **SRR6656087** |
| 85 | **SRR6656088** |
| 86 | **SRR8242276** |
| 87 | **SRR8242277** |
| 88 | **SRR8242278** |
| 89 | **SRR8242279** |
| 90 | **SRR8242280** |
| 91 | **SRR8242281** |
| 92 | **SRR8242282** |
| 93 | **SRR8242283** |
| 94 | **SRR8242284** |
| 95 | **SRR8242285** |
| 96 | **SRR8242286** |
| 97 | **SRR8242287** |
| 98 | **SRR8242288** |
| 99 | **SRR8242289** |
| 100 | **SRR8242290** |
| 101 | **SRR8242291** |
| 102 | **SRR8242292** |
| 103 | **SRR8242293** |
| 104 | **SRR8242294** |

| | |
|---|---|
| 105 | **SRR8242295** |
| 106 | **SRR8242296** |
| 107 | **SRR8242297** |
| 108 | **SRR8242298** |
| 109 | **SRR8242299** |
| 110 | **SRR9068902** |
| 111 | **SRR9068904** |
| 112 | **SRR9068905** |
| 113 | **SRR9068906** |
| 114 | **SRR9068907** |
| 115 | **SRR9068908** |
| 116 | **SRR9068909** |
| 117 | **SRR9068910** |
| 118 | **SRR9068911** |
| 119 | **SRR9068912** |
| 120 | **SRR9068913** |
| 121 | **SRR9068914** |
| 122 | **SRR9068915** |
| 123 | **SRR9068916** |
| 124 | **SRR9068917** |
| 125 | **SRR9068918** |
| 126 | **SRR9068919** |
| 127 | **SRR9068920** |
| 128 | **SRR9068921** |
| 129 | **SRR9068922** |
| 130 | **SRR9068923** |
| 131 | **SRR9068924** |
| 132 | **SRR9068925** |
| 133 | **SRR9068926** |
